# Cell-specific discrimination of desmosterol and desmosterol mimetics confers selective regulation of LXR and SREBP pathways in macrophages

**DOI:** 10.1101/263434

**Authors:** Evan D. Muse, Shan Yu, Chantle R. Edillor, Jenhan Tao, Nathanael J. Spann, Ty D. Troutman, Jason S. Seidman, Adam Henke, Jason T. Roland, Katherine A. Ozeki, Jeffrey G. McDonald, John Bahadorani, Sotirios Tsimikas, Tamar R. Grossman, Matthew S. Tremblay, Christopher K. Glass

**Author notes:** These authors contributed equally to this work as co-first authors (Evan D. Muse, Shan Yu). These authors contributed equally to this work as co-second authors (Chantle R. Edillor, Jenhan Tao). Corresponding Author: Christopher K. Glass, University of California, San Diego, Department of Cellular & Molecular Medicine, School of Medicine, GPL Rm 217, 9500 Gilman Drive, Mail code 0651, La Jolla, CA, 92093.

## Abstract

Activation of liver X receptors (LXRs) with synthetic agonists promotes reverse cholesterol transport and protects against atherosclerosis in mouse models. Most synthetic LXR agonists also cause marked hypertriglyceridemia by inducing the expression of SREBP1c and downstream genes that drive fatty acid biosynthesis. Recent studies demonstrated that desmosterol, an intermediate in the cholesterol biosynthetic pathway that suppresses SREBP processing by binding to SCAP, also binds and activates LXRs and is the most abundant LXR ligand in macrophage foam cells. Here, we explore the potential of increasing endogenous desmosterol production or mimicking its activity as a means of inducing LXR activity while simultaneously suppressing SREBP1c induced hypertriglyceridemia. Unexpectedly, while desmosterol strongly activated LXR target genes and suppressed SREBP pathways in mouse and human macrophages, it had almost no activity in mouse or human hepatocytes *in vitro*. We further demonstrate that sterol-based selective modulators of LXRs have biochemical and transcriptional properties predicted of desmosterol mimetics and selectively regulate LXR function in macrophages *in vitro* and *in vivo*. These studies thereby reveal cell-specific discrimination of endogenous and synthetic regulators of LXRs and SREBPs, providing a molecular basis for dissociation of LXR functions in macrophages from those in liver that lead to hypertriglyceridemia.

**SIGNIFICANCE STATEMENT:** The beneficial effects of LXR pathway activation in the prevention of atherosclerotic heart disease have long been appreciated. However, efforts to translate those effects in humans with synthetic LXR ligands has been met with the unintended consequence of hypertriglyceridemia, a product of co-activation of SREBP1c. Natural LXR ligands such as desmosterol do not promote hypertriglyceridemia because of coordinate down-regulation of the SREBP pathway. Here, we demonstrate that synthetic desmosterol mimetics activate LXR pathways macrophages both in vitro and in vivo without co-stimulation of SREBP1c. Unexpectedly, desmosterol and synthetic desmosterol mimetics almost no effect on LXR activity in hepatocytes in comparison to conventional synthetic LXR ligands. These findings reveal cell-specific differences in LXR responses to natural and synthetic ligands in macrophages and liver cells that provide a conceptually new basis for future drug development.

## INTRODUCTION

Although improvements in the prevention and treatment of cardiovascular disease (CVD) over the last decade have contributed to a significant reduction in the burden of CVD, it still accounts for over a third of all deaths in the United States and worldwide each year (*1*). In fact, more people die each year secondary to CVD than any other cause, with coronary heart disease and stroke representing the majority of cases (*2*). Increased apolipoprotein B-100 (ApoB)-associated lipid species, namely LDL cholesterol (LDL-C), remains one of the best-appreciated risk factors for atherosclerotic heart disease. Accordingly, reduction of LDL-C through the use of statins or recently developed antibodies directed against proprotein convertase subtilisin/kexin type 9 (PCSK9) represents one of the mainstays of preventive therapy (*3, 4*). However, myocardial infarction and stroke still occur in a subset of individuals despite cholesterol lowering, and therapies directed at additional targets are of potential clinical benefit.

Macrophages are key cellular players in the initiation and progression of atherosclerosis through their roles in uptake of modified lipoproteins in the artery wall, production of inflammatory mediators, and secretion of metalloproteases that contribute to plaque instability (*5-8*). A subset of macrophages within atherosclerotic lesions are characterized by massive accumulation of cholesterol esters in lipid droplets, resulting in a ‘foam cell’ phenotype indicative of a failure of normal cholesterol homeostasis. Given their central role in integrating both cholesterol homeostasis and inflammatory signaling in macrophages, the liver X-receptors (LXR) represent logical targets for pharmacologic intervention in atherosclerosis (*9-11*). LXR activation is known to promote cholesterol efflux in macrophages by activation of ATP-binding cassette transporter A1 and G1 (ABCA1/ ABCG1) (*12, 13*) while also repressing the pro-inflammatory products of NF-κB signaling (*14*). Stimulation of cholesterol efflux in macrophages and other cell types contributes to overall functions of LXRs in mediating reverse cholesterol transport from peripheral cells to the liver for biliary secretion (*15, 16*).

Consistent with these homeostatic functions, deletion of LXRs either at the whole body level or within the hematopoietic compartment results in accelerated atherosclerosis in mouse models (*17, 18*). Conversely, administration of potent synthetic LXR agonists, such as GW3965, inhibits the development of atherosclerosis in these models (*19-21*). However, most synthetic agonists of LXR have also been found to strongly activate SREBP1c and downstream genes involved in fatty acid biosynthesis, including fatty acid synthase (FAS), that subsequently lead to increased serum cholesterol and triglyceride levels (*22, 23*). Thus, while activating LXR has positive effects in the prevention of atherosclerosis in terms of enhancing reverse cholesterol transport and suppression of pro-inflammatory pathways, the negative aspect of hypertriglyceridemia and fatty liver – a product of SREBP activation – has prevented synthetic LXR agonists from being clinically useful therapeutics. Empiric efforts to develop ‘dissociated’ LXR agonists that retain the ability to activate LXRs but do not induce hypertriglyceridemia have been partially successful (*24-27*), but underlying mechanisms are poorly understood.

LXR activity is normally induced under conditions of cholesterol excess in a manner that is reciprocal to coordinate inhibition of the processing of the SREBP transcription factors. LXRs do not sense cholesterol directly, but are instead positively regulated by oxysterols and intermediates in the cholesterol biosynthetic pathway (*28-30*). In contrast to most synthetic LXR agonists, natural LXR agonists also suppress processing of the SREBP proteins (*31*). In the case of oxysterols, such as 25-hydroxy cholesterol, inhibition is mediated through interactions with the INSIG proteins that prevent trafficking of SREBPs to the Golgi for proteolytic activation (*32*). The cholesterol biosynthetic intermediate desmosterol was first noted to be an endogenous LXR-activating ligand in studies of plant sterols and sterol intermediates in the cholesterol biosynthetic pathway (*30*). In contrast to oxysterols, desmosterol most likely suppresses SREBP processing by interacting with SCAP to retain the SREBPs in the endoplasmic reticulum (*31*).

In a lipidomic analysis of murine macrophage foam cells and human atherosclerotic plaques, desmosterol was found to be the most abundant endogenous LXR activator (*33*). The accumulation of desmosterol in macrophage foam cells was correlated with downregulation of *Dhcr24*, which encodes the 24-dehydroxycholesterol reductase enzyme that converts desmosterol to cholesterol (**Figure 1A**). Notably, treatment of macrophages with increasing concentrations of desmosterol led to coordinate increases in LXR-dependent pathways and suppression of SREPB-pathways. As a consequence, genes involved in cholesterol efflux were induced, while genes involved in cholesterol and fatty acid synthesis were downregulated (*33*). These findings confirmed the prediction that desmosterol could balance lipid homeostasis via reciprocal actions on LXR and SREBP activities (*30*).

**Figure 1.**
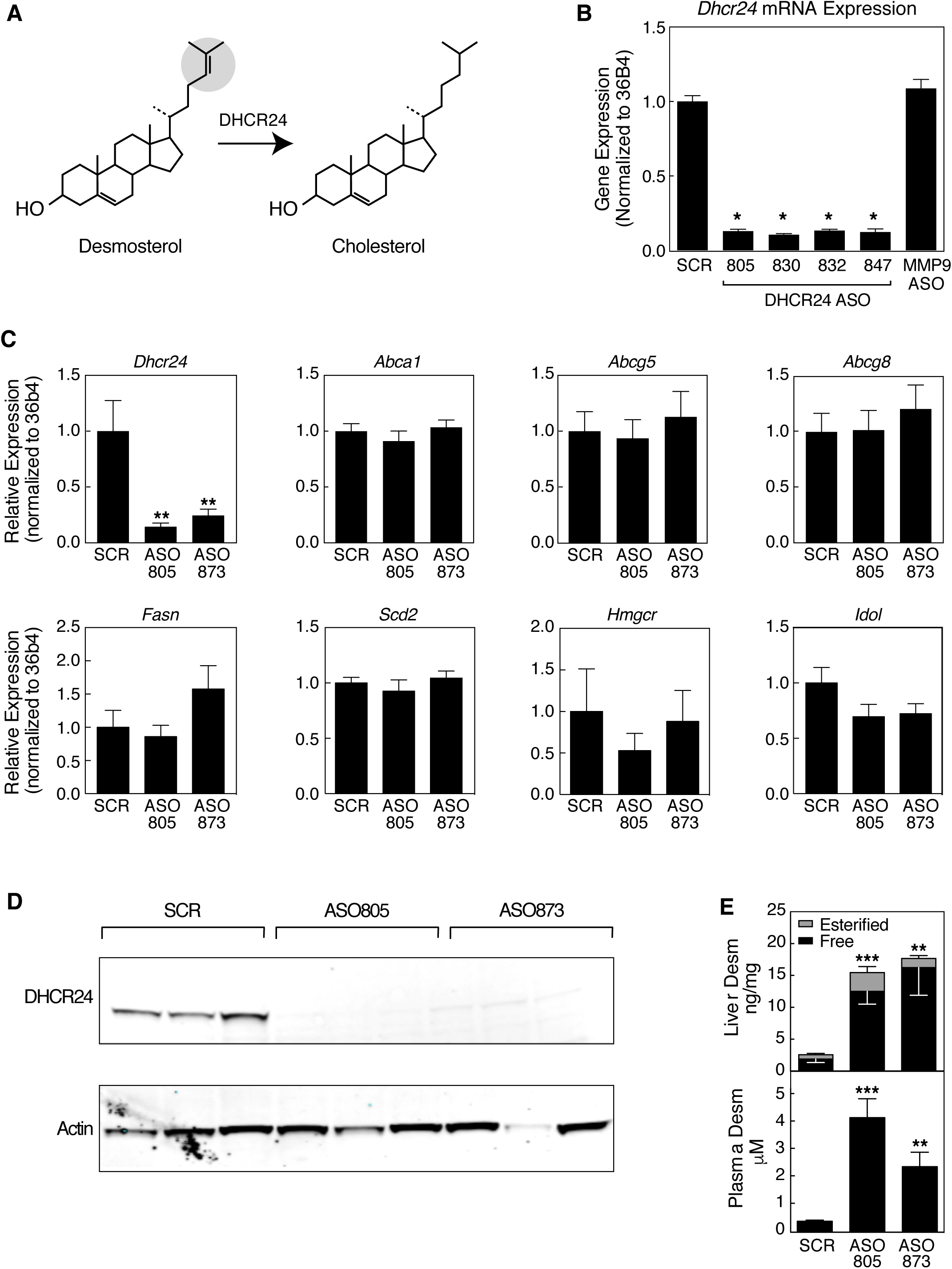
The effect of 24-dehydrocholesterol reductase (Dhcr24)-specific antisense oligonucleotide (ASO) treatment in mice. **A**. Catalysis of the final step in cholesterol biosynthesis by *Dhcr24.***B**. Effect of treatment of mouse thioglycolate elicited macrophages (TGEMs) with four separate ASOs to *Dhcr24* (ION-599805, ION-599830, ION-599832 and ION-599847, hereafter indicated as 805, 830, 832 and 847, respectively) or an ASO directed at *Mmp9* (MMP9 ASO), as assessed by RT-qPCR. (* p < 0.0001 vs SCR) **C**. Gene expression levels of the indicated genes in mouse liver after treatment with ASO. (** p < 0.001). **D.** Western immunoblot analysis of Dhcr24 and Actin in liver protein extracts in mice treated with the indicated *Dhcr24*-specific ASOs. **E**. Plasma and liver desmosterol concentrations as measured by LC-MS in mice treated with *Dhcr24*-specific as compared with SCR ASOs. (** p < 0.01, *** p < 0.001).

These observations raised the questions of whether the desmosterol pathway operates in other cell types and whether it would be possible to activate this pathway, or mimic it, as a therapeutic strategy. Studies in mouse macrophages pointed to downregulation of *Dhcr24* as being the key event leading to accumulation of desmosterol (*33*). One straightforward strategy would thus be to inhibit DHCR24 activity. Remarkably, this was first achieved more than 50 years ago following the identification of triparanol (also known as mer-29) as a potent inhibitor of DHCR24. Administration of triparanol to hypercholesterolemic human subjects led to marked reductions in serum cholesterol and corresponding rises in circulating desmosterol (*34, 35*). However, triparanol was rapidly withdrawn from the market after reports of severe cataracts and alopecia (*36-38*) and there is no evidence whether there was an impact on the development of cardiovascular diseases. While there are dermatologic manifestations of desmosterolosis, it is unknown if the extent of alopecia or development of cataracts is specifically related to increased circulating desmosterol or potentially an off-target effect of triparanol (*39*).

Here, we investigate the potential to modulate and mimic the desmosterol pathway *in vivo* and *in vitro* as a means of coordinately regulating the LXR and SREBP pathways. To modulate the desmosterol pathway *in vivo*, we developed potent and specific anti-sense oligonucleotides (ASOs) that reduce *Dhcr24* expression in liver by more than 80%. Unexpectedly, while this treatment resulted in significant increases in endogenous desmosterol, no significant changes in LXR or SREBP target genes were observed in the liver. To mimic the desmosterol pathway, we demonstrate that previously reported selective LXR modulators DMHCA (*27*) and MePipHCA (*40*) have properties of synthetic desmosterol mimetics. While desmosterol, DMHCA and MePipHCA coordinately regulate the LXR and SREBP pathways in primary mouse and human macrophages, they exhibited very little effect on gene expression in primary mouse and human hepatocytes. The differential effects of DMHCA and MePipHCA on LXR and SREBP target genes in macrophages and hepatocytes are also observed *in vivo*. Remarkably, LXR target genes are activated in Kupffer cells in response to DMHCA, in contrast to liver as a whole. In concert, these findings suggest a molecular basis for dissociation of LXR functions in macrophages and hepatocytes that would enable retention of anti-atherogenic properties without promoting hypertriglyceridemia.

## RESULTS

### Blockade of DHCR24 using gene-specific ASO leads to increased endogenous desmosterol without potentiating LXR-mediated target genes

To modulate the endogenous desmosterol pathway for coordinate regulation of LXR and SREBP target genes *in vivo*, we developed antisense oligonucleotides (ASOs) specific to 24-dehydrocholesterol reductase (DHCR24). Based on the effects of inhibition of DHCR24 by triparanol and genetic deficiency of *Dhcr24*, we hypothesized that reduction of *Dhcr24* expression would lead to increased desmosterol levels and corresponding changes in LXR- and SREBP-dependent gene expression (**Figure 1A**). Out of more than fifty potential *Dhcr24* specific ASOs developed and initially assayed (data not shown) we tested four of the most active ASOs in plated thioglycolate elicited macrophages (TGEMs) (**Figure 1B**). After 48 hours of exposure to ASO, *Dhcr24* gene expression as assessed through quantitative real-time polymerase chain reaction (qRT-PCR) was less than a quarter of scramble control (SCR) with four separate *Dhcr24*-specific ASOs (ION-599805, ION-599830, ION-599832 and ION-599847, hereafter referred to as 805, 830, 832 and 847, respectively). *Dhcr24* gene expression was also unchanged by a negative control MMP9 ASO compared to the SCR-treated cells. We then aimed to use these ASOs to reduce *Dhcr24* expression in C57Bl/6 mice.

We delivered control (SCR) ASO and two separate DHCR24-specific ASO to six mice per group via biweekly intraperitoneal (i.p.) injections over a three-week period (**Supplemental Figure 1A**). The treatment regimen was well tolerated in all study groups with expected weight gain and no difference in body weights at the termination of the study (**Supplemental Figure 1B**). Following three weeks of treatment, hepatic expression of *Dhcr24* was markedly reduced in both cohorts of *Dhcr24* ASO treated animals as compared with SCR control (15% and 25% vs. SCR control for 805 and 873, respectively, p < 0.001) (**Figure 1C**). This was mirrored by a substantial reduction in DHCR24 gene product as examined by Western immunoblot analysis of liver extract (**Figure 1D**). Concomitant with this reduction of *Dhcr24* gene expression and protein, circulating plasma desmosterol increased by 10-fold after treatment with *Dhcr24* ASO (0.38 ± 0.04 μM, 3.97 ± 0.65 μM, 2.27 ± 0.50 μM for SCR, 805 and 873, respectively) as measured by LC-MS (**Figure 1E**). In addition, hepatic desmosterol levels increased after *Dhcr24* ASO treatment (1.87 ± 0.53 ng/mg, 12.50 ± 2.01 ng/mg, 16.23 ± 4.34 ng/mg for SCR, 805 and 873, respectively) attributable mainly to increases in free rather than esterified desmosterol (**Figure 1E**). Surprisingly, despite these increases in circulating and hepatic desmosterol there were no observed alterations in the expression of the key hepatic LXR-target genes ATP-binding cassette transporter subfamily A, member 1 (*Abac1*), ATP-binding cassette transporter subfamily G, member 5 (*Abcg5*) or ATP-binding cassette transporter subfamily G, member 8 ( *Abcg8)* in *Dhcr24* ASO treated animals versus SCR control (**Figure 1C**). Nor were there differences in the hepatic expression of SREBP-target genes (**Figure 1C)**.

In a second experimental paradigm, we performed 1 week of subcutaneous (s.q.) ASO treatment in C57BL/6 mice who received thioglycolate 4 days prior to study end. Both liver and TGEMs showed a reduction in *Dhcr24* gene expression, although macrophages responded less robustly than liver (95% and 73% reduction in liver and macrophage of *Dhcr24* ASO vs SCR control treated animals, respectively) (**Supplemental Figure 1C**). As in the case of liver, the major LXR-target genes *Abca1* and *Abcg1* were not significantly modulated (p= ns for both, *Dhcr24* ASO vs SCR ASO) despite increased desmosterol concentrations in plasma, liver and macrophage **(Supplemental Figure 1D)**. Taken together, these data show that while *Dhcr24*-specific ASOs reduced hepatic and macrophage expression of *Dhcr24* mRNA and protein leading to increased levels of cellular and circulating desmosterol, they did not lead to activation of LXR-target genes or suppression of SREBP target genes.

### Cell-type specific effect of desmosterol in macrophage as compared to hepatocytes

The lack of an effect of knockdown of *Dhcr24* in liver on LXR- or SREBP target genes despite a significant increase in hepatic desmosterol raised the questions of whether desmosterol reached sufficient intracellular concentrations to be active and/or whether the desmosterol pathway is utilized in the liver. To directly compare responses of macrophages and hepatocytes to desmosterol, we evaluated the expression of *Dhcr24* and *Abca1* in plated TGEMs and *DHCR24* and *ABCA1* in HepG2 cells treated with increasing concentrations of desmosterol and the synthetic LXR-ligand T0901317 (**Figure 2A**). Whereas treatment with T0901317 resulted in an increase of *Abca1* expression in TGEMs (19.05 ± 3.86, p < 0.01 for 1 μM T0901317 relative to vehicle) and *ABCA1* expression in HepG2 (1.80 ± 0.30, p < 0.01 for 1 μM T0901317 relative to vehicle) cells, this LXR-target gene was up-regulated only in TGEMs (8.51 ± 1.40, p < 0.01 relative to vehicle) but not HepG2 cells after treatment with 10 μM desmosterol. In addition, while there was no effect of T0901317 on the suppression of the SREBP-target gene *Dhcr24* in either cell type, treatment with 10 μM desmosterol resulted in a marked down-regulation of *Dhcr24* (0.19 ± 0.08, p < 0.01 relative to vehicle) solely in TGEMs (**Figure 2A**).

**Figure 2.**
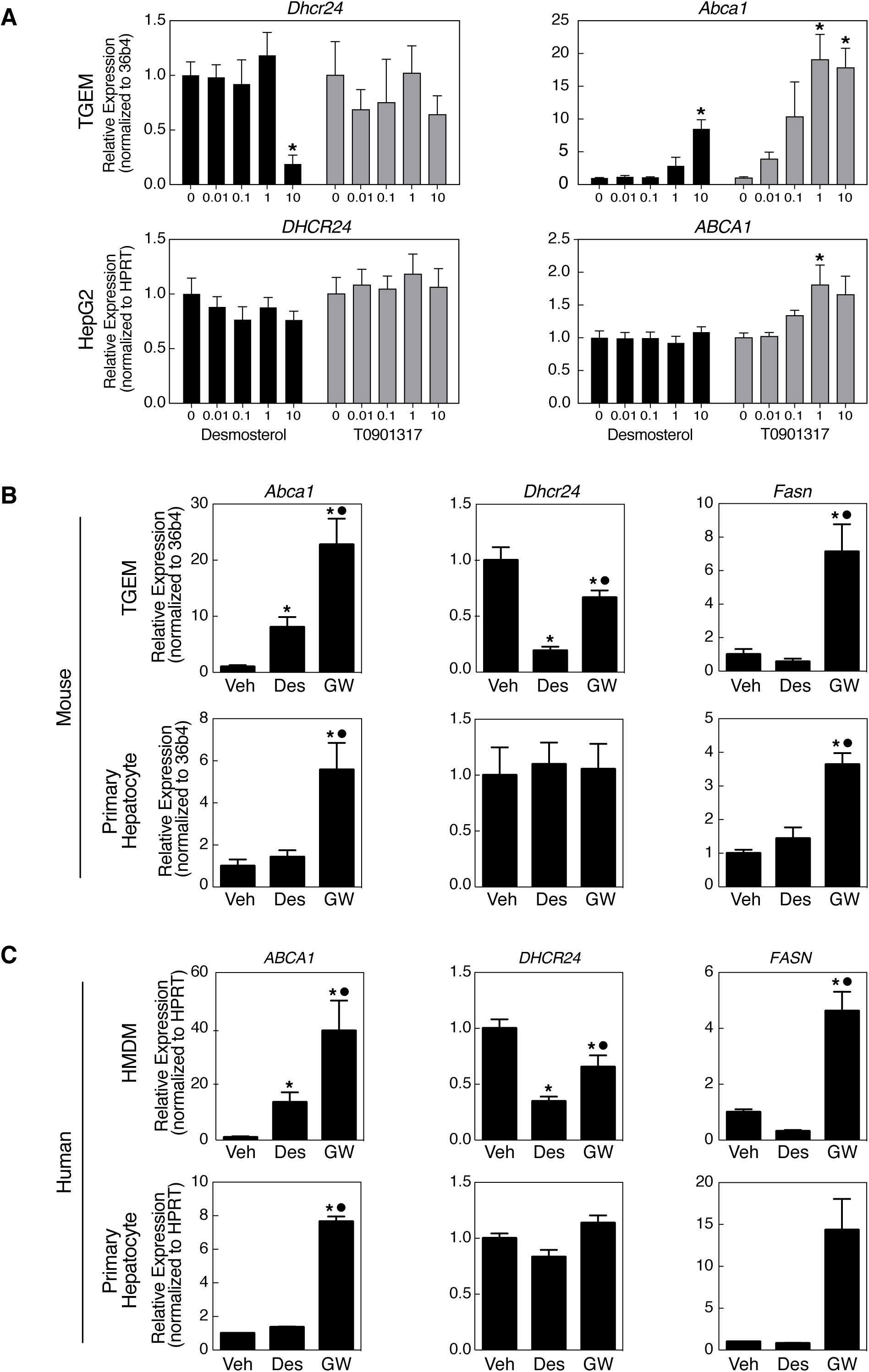
LXR- and SREBP-mediated gene expression profiles in mouse and human macrophages and hepatocytes. **A.** Dose dependent modulation of *Dhcr24* and *ABCA1*in plated TGEM and HepG2 cells by desmosterol and T0901317. (* p < 0.01). **B.** In mouse TGEM desmosterol (Des, 10 μM) induces expression of *Abca1* while suppressing *Dhcr24* without affecting *Fasn* whereas the synthetic LXR ligand GW3965 (GW, 1 μM) induces the expression of *Fasn*. In primary mouse hepatocytes, desmosterol fails to illicit changes in Abca1 and Dhcr24 while GW3965 retains its ability to activate LXR and SREBP. (* p < 0.05 vs Veh; • p < 0.05 vs desmosterol). **C**. The action of desmosterol (Des, 10 μM) and GW3965 (GW, 1 μM) on *ABCA1*, *DHCR24* and *FASN* in cultured human monocyte derived macrophages (HMDM) and human primary hepatocytes recapitulates the transcriptional activities observed in mouse TGEM and hepatocytes. (* p < 0.05 vs Veh; • p < 0.05 vs desmosterol).

To further explore the specific LXR- and SREBP-dependent gene responses to desmosterol we performed qRT-PCR analysis in primary macrophages and primary hepatocytes from both mice and humans ( **Figure 2B** and **2C**). Treatment of TGEMs with 10 μM desmosterol resulted in an increase in the LXR-responsive gene *Abca1* (8.06 ± 1.80, p < 0.05 relative to vehicle) and a decrease in *Dhcr24* (0.19 ± 0.04, p < 0.05 relative to vehicle) without subsequent activation of the SREBP-responsive gene fatty acid synthase (*Fasn*), whereas it had no effect in primary mouse hepatocytes (**Figure 2B**). In contrast, treatment of cells with the synthetic selective LXR ligand GW3965 resulted in increased levels of *Abca1* (22.77 ± 4.62, p < 0.05 relative to vehicle) and *Fasn* (7.13 ± 1.64, p < 0.05 relative to vehicle) mRNA in both macrophages and hepatocytes (5.58 ± 1.28, p < 0.05 and 3.64 ± 0.34, p < 0.05 for *Abca1* and *Fasn*, respectively) (**Figure 2B**). The LXR and SREBP-target gene expression changes induced by desmosterol and GW3965 in mouse macrophages and hepatocytes were fully recapitulated in human monocyte-derived macrophages (HMDM) and primary hepatocytes (**Figure 2C**). Collectively, these data suggest that while desmosterol effectively leads to a dose-dependent increase in LXR target genes without inducing SREBP-target genes as is seen for synthetic selective LXR ligands in both human and mouse macrophages, these effects are largely absent in hepatocytes.

### Synthetic desmosterol mimetics exhibit a cell-type specific LXR activation profile similar to desmosterol

Appreciating the differential LXR and SREBP target gene response in macrophages and hepatocytes of desmosterol as compared to conventional LXR ligands (e.g., GW3965, T0901317), we sought synthetic compounds that might represent desmosterol mimetics. Such molecules by definition would function as agonists of LXRs and antagonists of SREBP processing by interacting with SCAP. While the chemical structures of GW3965 and T0901317 are vastly dissimilar to desmosterol (**Figure 3A**), DMHCA, a previously reported synthetic dissociated LXR agonist with anti-atherosclerotic potential, retains much of the same desmosterol chemical backbone (24, 26, 27). In TGEMs, DMHCA not only activates LXR target genes such as *Abca1* and fails to induce *Fasn* expression, as previously reported, but also strongly represses the SREBP target gene *Dhcr24* (**Figure 3B**). This activity profile is thus consistent with that of a desmosterol mimetic. In addition, we evaluated recently developed derivatives of DMHCA, exemplified by MePipHCA, which we recently reported as a dissociated LXR agonist that inhibits inflammation in models of traumatic brain injury and inflammatory bowel disease (41). Similar to DMHCA, MePipHCA not only activated the LXR target genes in macrophages, but also strongly suppressed *Dhcr24* expression (**Figure 3C**). We also observed that, in contrast to T0901317, DMHCA and MePipHCA did not cause lipid accumulation in HepG2 cells after 72 hour treatment (**Figure 3D**), consistent with a lack of an effect of DMHCA and MePipHCA on SREBP1c expression and lipid biogenesis.

**Figure 3.**
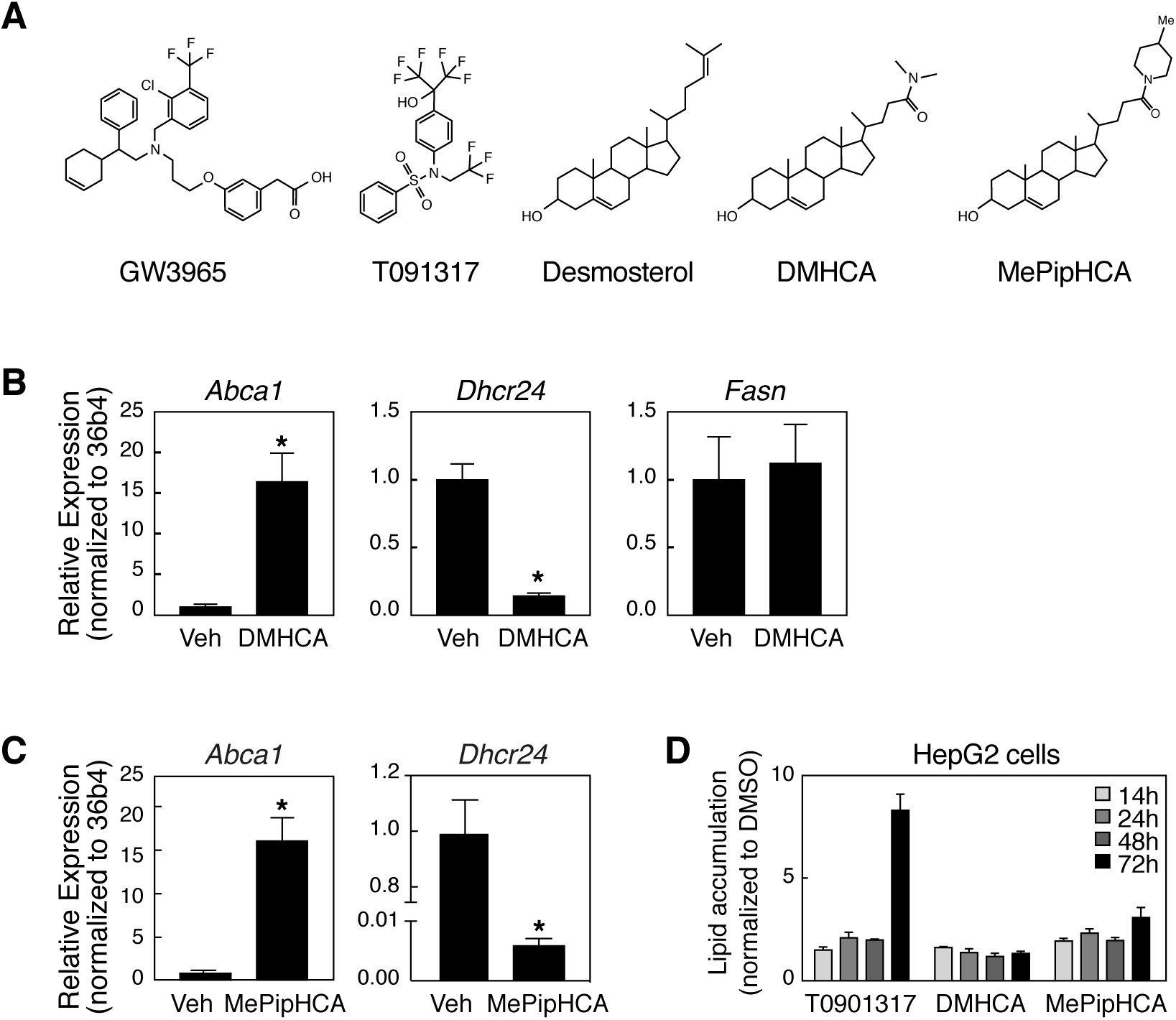
DMHCA and MePipHCA have desmosterol-like activities. **A.** Comparison of the chemical structures of GW3965, T0901317, desmosterol, DMCHA and MePipHCA. **B.** Effects of DMHCA (1 μM) on expression of *Abca1, Dhcr24* and *Fasn* in mouse TGEM. (* p < 0.01 vs Veh). **C.** Effects of MePipHCA (1 μM) on *Abca1* and *Dhcr24* expression in mouse TGEM. (* p <0.01 vs Veh). **D.** Effects of T0901317, DMHCA and MePIPHCA on lipid accumulation in HepG2 cells.

### Whole transcriptome assessment of desmosterol mimetics

Having appreciated the cell-type specific characteristics of desmosterol signaling, the similarities shared with DMHCA, especially in terms of the dissociation of LXR and SREBP pathway regulation, and promise of the structurally related mimetic MePipHCA, we conducted an unbiased, whole transcriptome comparison of these ligands with the conventional LXR-ligands GW3965 and T0901317 using RNA-sequencing (RNA-Seq) analysis. We began by treating plated TGEM and mouse primary hepatocytes with desmosterol (10 μM), DMHCA (1 μM), MePipHCA (1 μM), GW3965 (1 μM) and T0901317 (1 μM). Representative scatter plots show that while treatment with desmosterol, DMHCA or T0901317 leads to a robust LXR response in TGEMs, the overall LXR response is absent in hepatocytes except after treatment with T0901317 (**Figure 4A)**. Heat maps of individual genes from key LXR and SREBP pathways illustrate that while all compounds tested in TGEMs induce *Abca1* expression (LXR-responsive gene), desmosterol, DMHCA and MePipHCA repress SREBP-responsive genes (*Dhcr24*, *Hmgcr* and *Ldlr*) while GW3965 and T0901317 activate *Fasn* and *Srebf1* (**Figure 4B**). In mouse primary hepatocytes, only GW3965 and T0901317 retain the expected LXR activities and activate SREBP1c target genes such as *Fasn*. Gene ontology (GO) term analysis of specific up- and down-regulated gene pathways reinforced the LXR-activating effect of all compounds in TGEMs and the combined LXR and SREBP1c activating effect of GW3965 and T0901317 in both TGEMs and hepatocytes (**Figure 4C**). Notably, the SREBP-mediated pathways noted by the GO terms “Cholesterol metabolic process” and “Sterol biosynthetic process” are markedly suppressed by desmosterol, DHMCA and MePipHCA in macrophages (**Figure 4C**). In principle component analysis (PCA) performed on the differentially expressed genes (up and down) in both TGEMs (left panels) and primary hepatocytes (right panels), the conventional ligands GW3965 and T0901317 cluster tightly together, as do desmosterol and DMHCA, with MePipHCA assorting itself distinctly along the first two eigenvectors (**Figure 4D**). The divergence of MePipHCA for upregulated genes is partly driven by its activation of several genes that have functional annotations associated with the “Response to unfolded protein” pathway in GO term analysis. Of note, this pathway is also activated by 25-hydroxy cholesterol in macrophages (42).

**Figure 4.**
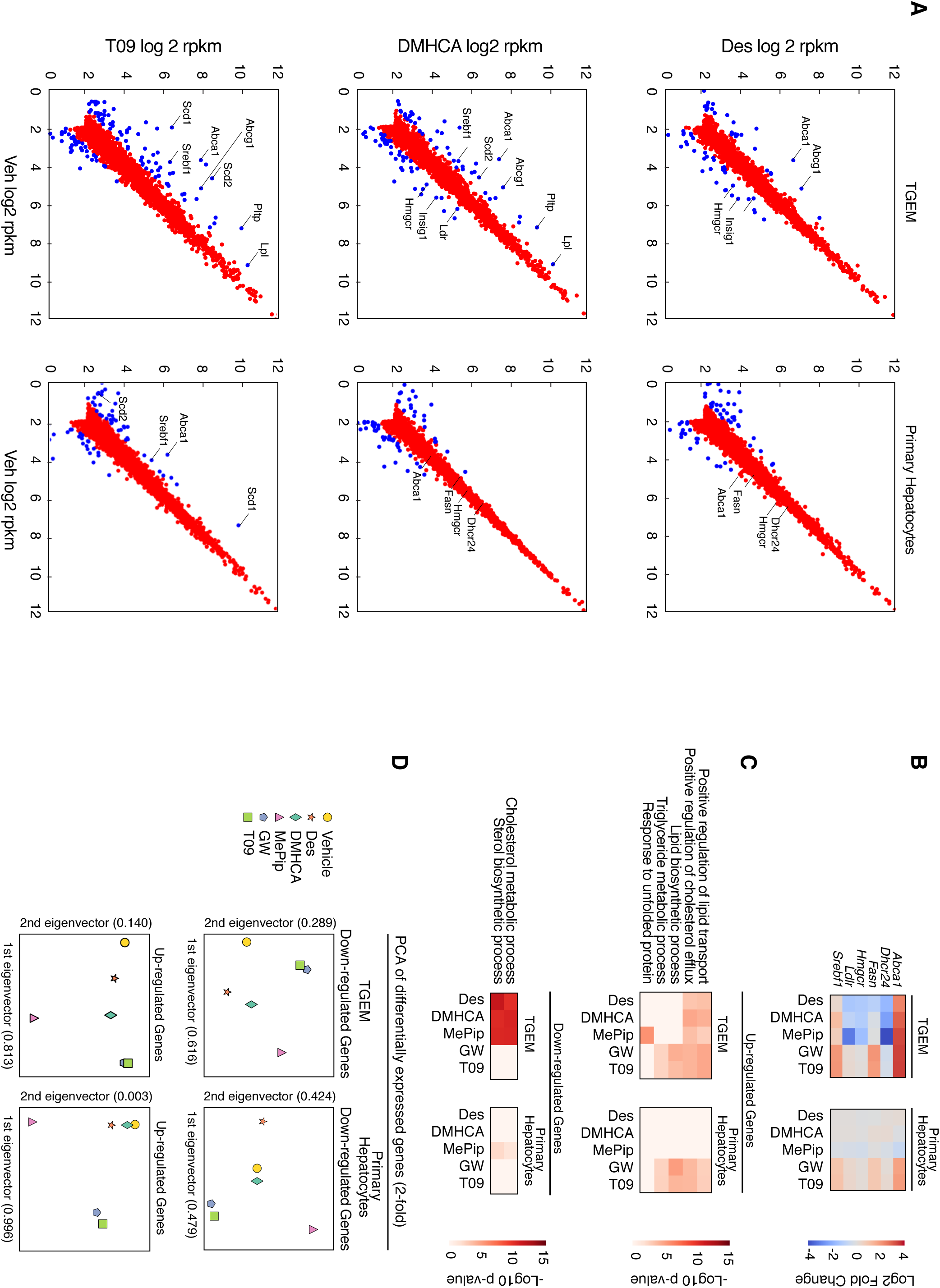
Whole transcriptome RNA-sequencing assessment of mouse TGEM and primary hepatocytes. **A.** Scatter plots for up and downregulated genes in TGEM (n = 3 per treatment) and primary hepatocytes (n = 2 per treatment) after exposure to desmosterol (10 μM), DMHCA (1 μM) and T0901317 (1 μM). Key LXR- and SREBP-target genes are highlighted. Genes noted in blue have 2-fold changes versus vehicle with FDR < 0.05. **B.** Heat map for key genes illustrating Log2-fold change compared to vehicle in TGEM and primary hepatocytes. **C.** GO-term analysis of up-regulated and down-regulated genes in response to various ligands. **D.** PCA analysis for all differentially expressed genes with 2-fold changes and FDR < 0.05 versus vehicle.

We observed similar differences in the transcriptional responses of macrophages and hepatocytes in our unbiased, whole transcriptome RNA-Seq analysis of human cells. First we compared the global response of up- and down-regulated genes in response to these ligands in both HMDM and human primary hepatocytes (**Figure 5A**). Notably, and similar to our studies in mice, whereas desmosterol, DMHCA and T0901317 initiate an overall program of gene activation and repression in HMDM, this response is blunted in primary hepatocytes treated with desmosterol and DMHCA but not those treated with T0901317. The most significant transcriptional pathways up- and down-regulated by these treatments in HMDM and human primary hepatocytes is relatively unchanged from mouse, with the coordinated activation of reverse cholesterol transport and suppression of cholesterol and triglyceride biosynthetic pathways observed in macrophage (**Figure 5B**). This pathway analysis is supported on the individual gene level (**Figure 5C**). In addition, HMDM from five individual patients with varying degrees of coronary atherosclerosis and comorbid chronic diseases were treated with desmosterol, DMHCA and T0901317. While the range of transcriptional responses to these ligands likely highlights the influence of natural genetic variation, all HMDMs showed robust increases in LXR-mediated genes (*ABCA1, ABCG1*) when treated with desmosterol, DMHCA and T0901317. T0901317 activated SREBP target genes (*SCD, FASN*), while these genes were suppressed or unchanged in response to desmosterol or DMHCA, respectively. Most apparent here is the global similarity of DMHCA with the natural oxysterol desmosterol in regards to LXR and SREBP transcriptional control, and their differences with the conventional LXR agonist T0901317 (**Figure 5C**). While desmosterol, DMHCA and MePipHCA activate LXR-mediated pathways and suppress SREBP-mediated pathways in macrophage of mice and humans, this response is absent or blunted in primary hepatocytes. Conversely, GW3965 and T0901317 remain potent activators of LXR-pathway genes in both cell types while activating SREBP-responsive genes in tandem.

**Figure 5.**
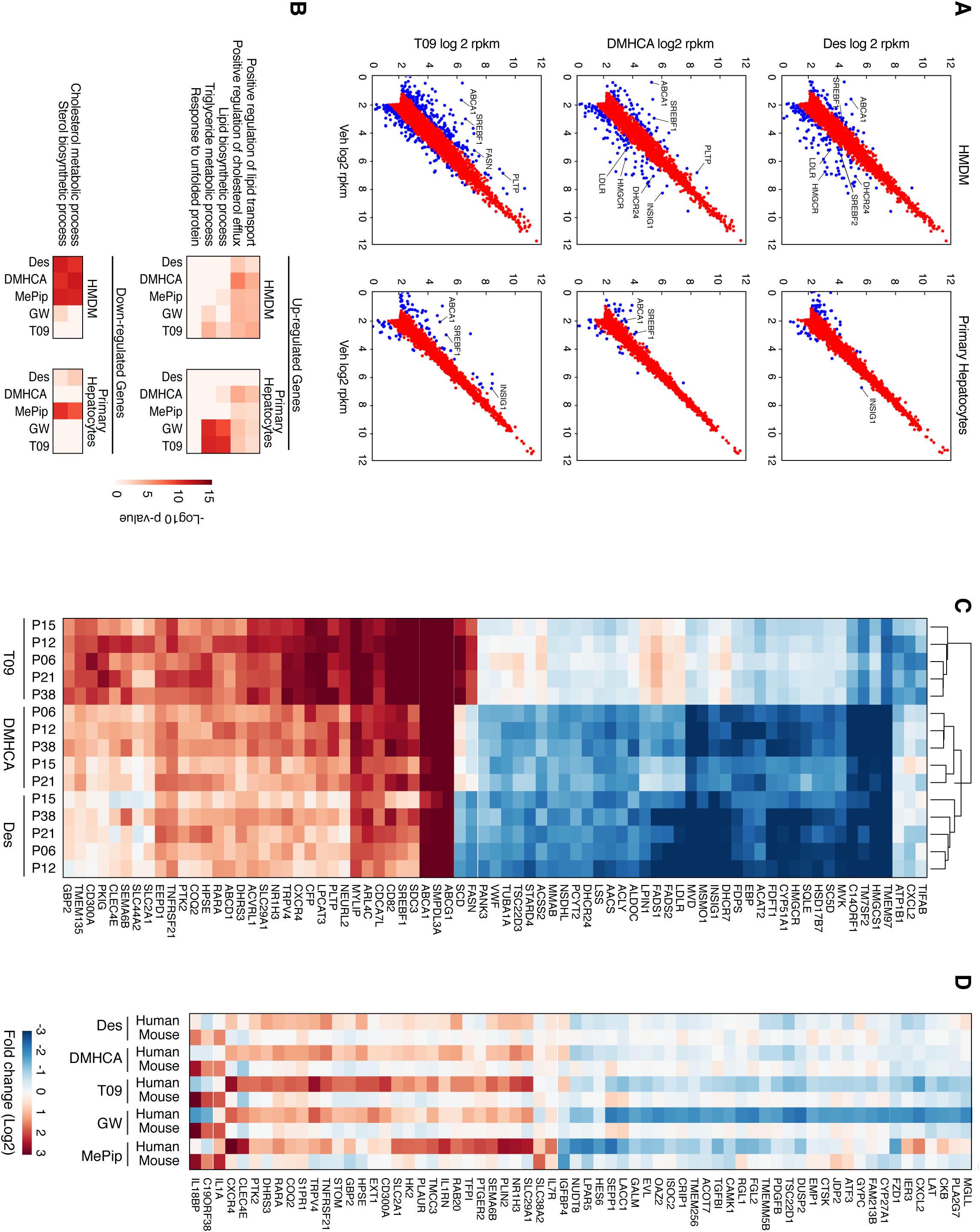
RNA-Seq analysis reveals an overall conserved transcriptional response to LXR ligands in human macrophage and primary hepatocytes as in mice however key differences remain. **A**. Scatter plots for up and downregulated genes in HMDM and primary hepatocytes after exposure to desmosterol (10 μM), DMHCA (1 μM) and T0901317 (1 μM). **B**. GO-term analysis of up-regulated and down-regulated genes in response to various ligands. **C**. Heat map illustrating a shared global gene expression pattern in individual HMDM (n = 5) in response to desmosterol and DMHCA that is divergent from T0901317. **D**. Heat map of genes that are differentially regulated in macrophage of mouse and humans.

While the vast majority of genes that were up- and down-regulated by desmosterol, DMHCA and T0901317 were shared in mouse and human macrophages, we also noted certain genes that were differentially regulated (**Figure 5D**). While the roles of many of these genes in lipid homeostasis and regulation of inflammation remain unappreciated, we did observe differences in key genes such as the nuclear receptors LXRα (*NR1H3*) and retinoic acid receptor (*RARA*), both differentially induced in HMDM, and several regulators of lipid metabolism such as fatty acid desaturase (*FADS1*), 7-dehydrocholesterol reductase (*DHCR7*) and farnesyl-diphosphate farnyltransferase (*FDFT1*), all suppressed in HMDM compared to mouse TGEM.

### The macrophage-specific induction of LXR-response genes without potentiation of SREBP pathways with synthetic desmosterol mimetics is recapitulated in vivo

We then sought to examine if the cell-type specific effects observed in plated macrophages and hepatocytes in response to desmosterol and desmosterol mimetics (DMHCA and MePipHCA) were recapitulated in an *in vivo* model. We treated mice with thioglycollate by i.p. injection four days prior to treatment with compounds in order to elicit macrophage accumulation in the peritoneum. DMHCA, MePipHCA and T0901317 (50 mg/kg) were then given by i.p. injection 6 hours and 16 hours before peritoneal macrophages and whole liver were isolated and prepared for gene expression analysis. Gene expression of the LXR-responsive gene *Abca1* was robustly induced in peritoneal macrophages at 6 hours by T0901317, DMHCA and MePipHCA, whereas only T0901317 induced a coordinate response in *Abcg5* in liver (**Figure 6A**). The LXR-activating effect for DMHCA and MePipHCA was attenuated at 16 hours in macrophages, though remained strong for T0901317 in both tissues. Mirroring what we had observed in *in vitro* studies, treatment with T0901317 lead to the induction of SREBP-responsive genes, illustrated here by *Fasn*, in both macrophages and liver, while DMHCA and MePipHCA did not induce *Fasn* in either tissue.

**Figure 6.**
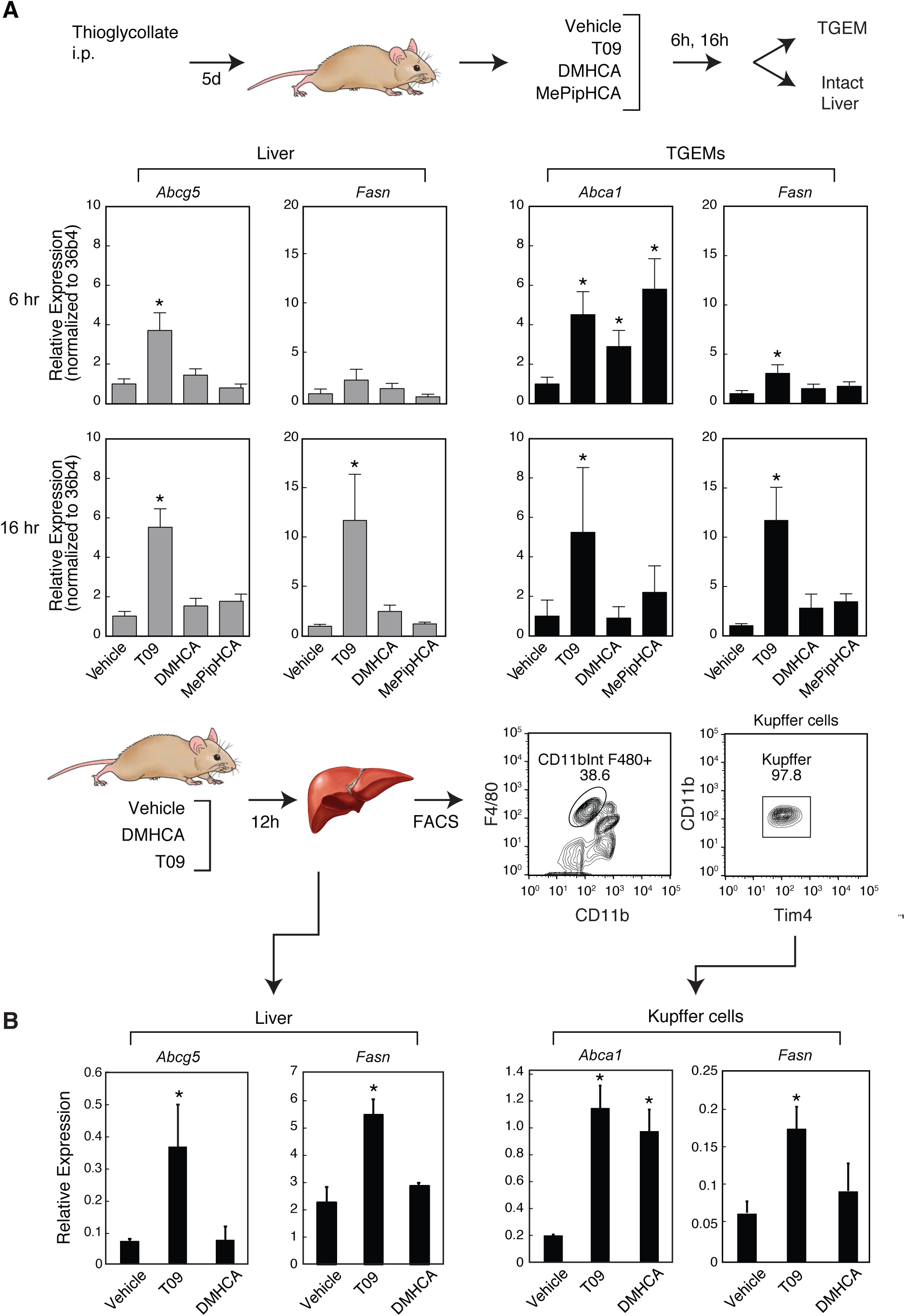
Cell-type specific effects on LXR and SREBP responsive genes in vivo. **A**. Gene expression profiling of LXR- and SREBP-target genes in mouse liver and peritoneal macrophages (TGEM) 6 and 16 hours after intraperitoneal (i.p.) administration of T0901317 (T09), DMHCA and MePipHCA as compared with vehicle (M-pyrol) (* p < 0.05 vs Veh). **B**. Gene expression profiles of *Abca1*, *Abcg5* and *Fasn* of isolated Kupffer cells and whole liver in mice treated with i.p. DMHCA, T0901317 (T09) or vehicle for 12h (* p < 0.05 vs Veh).

Although responses to DMHCA and MePipHCA were minimal or absent in intact liver, it was of interest to determine whether Kupffer cells, the resident macrophage population of the liver, exhibited similar responses to those observed in elicited peritoneal macrophages. To address this question, we treated mice with vehicle, DMHCA or T0901317 by i.p. injection, and 12h following injection Kupffer cells were purified by fluorescence activated cell sorting with target cells identified by CD45^+^F4/80^+^CD11b^int^Tim4^+^CD146^−^CD31^−^. As in the case of peritoneal macrophages, DMHCA was shown to selectively activate LXR-responsive genes in Kupffer cells (*Abca1*) but not whole liver (*Abcg5*) (**Figure 6B**). Whereas DMHCA had no effect on SREBP-responsive genes in either tissue compared to the dual activation of LXR- and SREBP-responsive genes by T0901317. These results indicate that these sterol-based synthetic LXR agonists (DMHCA and MePipHCA) appear to act preferentially in macrophages (and Kupffer cells) as compared to hepatocytes *in vivo* and, unlike T0901317, without potentiating SREBP1-responsive genes.

## DISCUSSION

Despite their key roles at the intersection of lipid metabolism and inflammation, the promise of LXRs as pharmacologic targets for the prevention and treatment of atherosclerotic heart disease has been limited by the difficulty of decoupling beneficial effects from activation of SREBP1-dependent pathways. Development of selective LXR modulators has been challenging in part because there are few evident rationale approaches for achieving this goal. A leading strategy has been to synthesize molecules that preferentially activate LXRβ, based on evidence for a dominant role of LXRα in driving SREBP1c expression and fatty acid biosynthesis in mouse liver (*9, 19, 41, 42*). However, a recent evaluation of a prototypic LXRβ-selective synthetic ligand demonstrated that in addition to induction of LXR target genes in human blood cells and inhibition of atherosclerosis in mouse models, it retained the ability to increase circulating triglyceride levels and hepatic triglycerides in human subjects (*43*).

Here, we have explored an alternative strategy that is based on the observation that most or all physiologic LXR agonists are also inhibitors of SREBP processing, either by binding to INSIGs (e.g., oxysterols) or SCAP (desmosterol). In contrast to synthetic agonists that selectively target LXRs, such endogenous ligands would be expected to activate genes involved in reduction of cellular cholesterol but have an attenuated effect on fatty acid biosynthesis due to inhibitory effects on processing of SREBP1c. We thus sought to test the hypothesis that raising endogenous levels of LXR agonists or mimicking their activity would have these effects.

Our initial *in vivo* approach was to increase intracellular concentrations of desmosterol by specifically reducing the activity of DHCR24 using antisense oligonucleotide technology. Unexpectedly, despite a marked elevation in hepatic and circulating desmosterol levels after treatment with *Dhcr24*-specific ASOs, we failed to observe concomitant activation of key LXR-response genes in liver or macrophages. One possible interpretation of this result is that, while significantly elevated, desmosterol did not reach intracellular concentrations required to activate LXRs. This hypothesis was supported by our observation that 10 μM desmosterol significantly elevated LXR-target genes in TGEM but not 1 μM desmosterol (**Figure 2A**). In addition, serum desmosterol levels of *Dhcr24* ASO treated mice were less than 20% of those observed in human subjects treated with triparanol or in the rare genetic disease of desmosterolosis (*39, 44, 45*). Additionally, although our studies were short term (1-3 weeks), we did not observe evidence of alopecia characteristic of *Dhcr24* knockout mice and humans treated with triparanol in the animals treated with *Dhcr24* ASO (unpublished observation). We conclude that raising endogenous desmosterol levels by reducing *Dhcr24* expression using ASOs is unlikely to be an effective strategy for activation of LXRs *in vivo*.

Given the inability to modulate the LXR axis by increasing endogenous desmosterol concentrations with *Dhcr24* ASO, we sought to specifically assess the desmosterol pathway in hepatocytes. Unexpectedly, we observed that concentrations of desmosterol that effectively activated LXR-responsive genes and suppressed SREBP-responsive genes in macrophages had little or no effect on these genes in mouse and human hepatocytes. Thus, these studies provide evidence for a cell-autonomous mechanism enabling cell-specific discrimination of an endogenous LXR ligand that confers selective activation of LXR target genes in macrophages.

Based on these findings, we characterized DMHCA, an empirically developed LXR agonist that exhibits anti-atherosclerotic activity without causing substantial hypertriglyceridemia in mice (*24*). Importantly, unlike conventional LXR agonists such as GW3965 and T090137, DMHCA is structurally related to desmosterol, raising the possibility that it functions as a desmosterol mimetic. Consistent with this possibility, DMHCA coordinately induced LXR target genes and repressed SREBP target genes. We therefore evaluated a series of derivatives capable of activating LXRs in transient transfection assays, the most potent of which was MePipHCA. This result provides evidence of the possibility to improve the physicochemical properties of this class of compounds for pharmaceutical use.

Genome-wide comparisons of desmosterol, DMHCA, MePipHCA, GW3965 and T090137 in mouse and human macrophages and hepatocytes strongly support the preferential activities of desmosterol and desmosterol mimetics in macrophages and demonstrate that they coordinately regulate the LXR and SREBP pathways in these cells. Desmosterol, DMHCA and MePipHCA regulated a highly overlapping set of genes, with DMHCA and MePipHCA exhibiting greater potency. Comparisons of the responses of mouse and human macrophages also indicated a high degree of similarity at the level of genes involved in cholesterol homeostasis. A relatively small number of genes exhibited species specific differences in responses, primarily in human monocyte derived macrophages, which are at this point of uncertain significance. There was relatively little individual variation in responses of the core set of LXR target genes involved in cholesterol homeostasis among the small set of human monocyte derived macrophages that were evaluated. Importantly, the macrophage-selective activities of DMHCA and MePipHCA were observed *in vivo*.

In addition to regulating the LXR/SREBP pathways, desmosterol was previously shown to inhibit inflammatory responses in macrophages, consistent with the actions of other LXR agonists (*33*). Although not evaluated for anti-inflammatory effects in these studies, we recently reported that MePipHCA significantly reduced disease severity and inflammatory markers in models of inflammatory bowel disease and traumatic brain injury without causing lipid accumulation in liver (*40*). A limitation of the current synthetic desmosterol mimetics is a relatively poor pharmacokinetic profile, requiring large doses for *in vivo* efficacy. Therefore, it is likely that substantial additional effort will be required to develop more drug-like molecules.

A major new and exciting question is the basis for cell-specific discrimination of desmosterol that confers preferential activity in macrophages. It is unlikely to be simple conversion of desmosterol to cholesterol or another LXR agonist because similar activities were observed for the synthetic agonists DMHCA and MePipHCA. A cell autonomous basis for this discrimination is strongly supported by the finding that Kupffer cells in the liver robustly respond to DMHCA, while surrounding hepatocytes do not. The mechanism presumably distinguishes between desmosterol/desmosterol mimetics and oxysterols, given the genetic evidence that 24 hydroxycholesterol, 25 hydroxycholesterol and 27 hydroxycholesterol are endogenous agonists of LXRs in the liver (*46*). We speculate that proteins involved in the intracellular transport of desmosterol/desmosterol mimetics restrict their access to SCAP and SREBPs in hepatocytes but not macrophages. Further understanding of the mechanism underlying differential actions of desmosterol in macrophages and hepatocytes thus remains an important future goal.

## MATERIALS & METHODS

### Reagents

Desmosterol was purchased from Avanti Polar Lipids (Alabaster, AL). DMHCA was resynthesized internally, along with the design and synthesis of MePipHCA. Mevastatin, mevalonolactone and M-pyrol were purchased from Sigma-Aldrich (St. Louis, MO). GW3965 and T0901317 were purchased from Cayman Chemicals (Ann Arbor, MI).

### Antisense Oligonucleotides

All antisense oligonucleotides (ASO) used for these studies were designed by Ionis Pharmaceuticals (Carlsbad, CA) to hybridize to the sequence spanning mouse 24-dehydrocholesterol reductase (DHCR24) mRNA. The scrambled control ASO is a chemistry control ASO that has the same length and chemical makeup as the DHCR24-specific ASO and is not expected to hybridize to any mRNA sequence. For *in vitro* cell studies ASOs were transfected into cells using Cytofectin (Genlantis, San Diego, CA) at 50 nM final concentration. For animal studies, ASOs were delivered in sterile saline at either 20 mg ASO per kg animal weight by twice weekly intraperitoneal (i.p.) injections or at 100 mg/kg delivered subcutaneously.

### Animals

Adult male C57BL/6 mice were acquired form Charles River Laboratories (Wilmington, MA). Mice were maintained in an IACUC-approved animal facility with a standard light-dark cycle and fed standard lab chow.

### Thioglycollate-Elicited Macrophage Generation

Peritoneal macrophages were harvested 4 days after intraperitoneal (i.p.) injection of thioglycollate. (Refer to supplement for additional details).

### Cell Culture

Thioglycollate-elicited macrophages (TGEMs) and HepG2 cells were maintained in RPMI-1640 with 10% FBS and 100 U/mL penicillin/streptomycin (Cellgro/Corning, Manassas, VA). Primary mouse hepatocytes were prepared as described and maintained in Hepatozyme medium with 10% FBS, 1% L-glutamine and 1% Penicillin Streptomycin. (Refer to supplement for additional details).

### Isolation and Culture of Human Monocyte Derived Macrophage

Human macrophages were prepared from CD14+ PBMCs isolated from human blood by incubating in RPMI-1640 + 10% FBS supplemented with penicillin/streptomycin and 50 ng/mL recombinant human macrophage colony stimulating factor (M-CSF) (R&D Systems, Minneapolis, MN). (Refer to supplement for additional details).

### In vivo injection of LXR agonists and purification of Kupffer cells

Mice were treated by i.p. injection with 50 mg/kg of DMHCA or T0901317 dissolved in 50:50 ethanol and M-pyrol at a concentration of 50 mg/ml. 12 hours later, mice were humanely euthanized by exposure to CO_2_ and whole liver pieces saved and liver non-parenchymal cells processed for fluorescence activated cell sorting of Kupffer cells, with modifications from published methodology (*47, 48*). (Refer to supplement for additional details).

### RNA Isolation and quantitative polymerase chain reaction

Total RNA was isolated with Trizol Reagent (Life Technologies) and Direct-zol RNA Spin Columns (Zymo Research, Irvine, CA). Total RNA was used for either first strand cDNA synthesis with SuperScriptIII (Life Technologies) or for RNA-sequencing (RNA-seq) library preparation. Real-time PCR reactions were prepared in 96-well plates using iTaq SYBR Green Supermix (BioRad, Hercules, CA) and performed on the StepOnePlus Quantitative Polymerase Chain Reaction Platform (Life Technologies) (Refer to supplement for additional details).

### RNA-Sequencing Library Preparation

Please see supplement for details. Briefly, polyA RNA was used for first-strand cDNA synthesis. Second strand synthesis was carried out using deoxy-UTP. Follwing end-repair the product was then incubated with EDAC SeraMag SpeedBeads and eluted with EB (Zymo). This was followed by dA-tailing, unique barcode adapter ligation and second strand digestion with UDG (Enzymatics). PCR amplification was performed by 12-15 cycles and size selected for 223-375 bp after separating on a 10% TBE gel (Life Technologies). The library was purified from the gel slice and single-end sequenced on a HiSeq 2000 instrument (Illumina, San Diego, CA).

### Plasma Analysis and Lipid Measurements

Plasma, liver and TGEMs were processed at the University of Texas Southwestern Medical Center for oxysterol and lipid metabolite analysis by liquid chromatography-mass spectrometry (LC-MS) as previously described in full: http://www.lipidmaps.org/protocols/index.html.

### Western Blot Analyses

Protein extracts were fractionated using a 4-12% Bis-Tris NuPAGE (Invitrogen, Carlsbad, CA, USA) gel and blotted onto Immobilon-P PVDF membrane (EMD Millipore, Billerica, MA). Membranes were incubated in primary antibody against mouse DHCR24 (Cell Signaling Technologies, Danvers, MA) or beta-actin (Santa Cruz Biotechnology, Dallas, TX) overnight at 4C) followed by secondary antibody conjugated to Alexa Fluor 680 (Molecular Probes, Eugene, OR) or IR Dye 800 (Rockland, Gilbertsville, PA). Membranes were then imaged using the Li-Cor Odyssey (Lincoln, NE). Please see supplement for details.

### LXRE Luciferase Activity Assay

Cells were transfected with lenti LXRα responding element luciferase reporter according to manufacturer’s instruction. The cells were incubated with compounds at indicated concentrations and the reporter gene activity signal was read by a PerkinElmer EnVision Multilabel Reader. Please see supplement for details.

### HCS LipidTOX neutral lipid stain assay

HepG2 cells were plated in 384 well plate at density of 5000 cells/well. The cells were treated with compounds at 1 μM concentration for different time points as indicated, followed by fixing with 3% formaldehyde and staining with HCS LipidTOX neutral lipid stain (Cat No: H34475, Invitrogen) according to the manufacturer’s protocol. The lipid content was quantified and analyzed by Cellomics high-content imaging system and software.

### Data Analysis

Experimental values are presented as the means of replicate experiments +/-standard error. Aside from the RNA-Seq experiments, comparisons among separate groups were made using the analysis of variance followed as well as the unpaired student’s t test with correction for multiple comparisons using Prism 7.0 software (GraphPad, La Jolla, CA). Statistical significance was defined as p < 0.05. Please see supplement for details regarding RNA-Seq analysis.

## Acknowledgements

We appreciate the generosity of the research volunteers who made parts of this study possible. We thank Leslie Van Ael for assistance with preparation of figures and Bonne Thompson for assistance with mass spectrometry analysis of sterols and oxysterols. These studies were supported by NIH grants HL088093, GM085764, DK063491 to CKG. CKG is also supported by the Ben and Wanda Hildyard Chair in Hereditary Diseases. EDM is supported by NIH/NCATS Clinical Translational Science Award 5KL2TR001112 and 5UL1TR001114 to Scripps Translational Science Institute. SY is supported by Crohn’s and Colitis Foundation of America Research Fellowship Award #368561. TDT was supported by the National Cancer Institute of the National Institutes of Health under Award Number T32CA009523. JSS was supported by an American Heart Association Fellowship 16PRE30980030 and NIH pre-doctoral training grant 5T32DK007541.

## Accession Number

Sequencing data supporting these studies can be found at the Gene Expression Omnibus database under accession numbers GSE90615.

## REFERENCES

1. D. Mozaffarian, E. J. Benjamin, A. S. Go, D. K. Arnett, M. J. Blaha, M. Cushman, S. R. Das, S. de Ferranti, J.-P. Després, H. J. Fullerton, V. J. Howard, M. D. Huffman, C. R. Isasi, M. C. Jiménez, S. E. Judd, B.M. Kissela, J. H. Lichtman, L. D. Lisabeth, S. Liu, R. H. Mackey, D. J. Magid, D. K. McGuire, E. R. Mohler, C. S. Moy, P. Muntner, M. E. Mussolino, K. Nasir, R. W. Neumar, G. Nichol, L. Palaniappan, D. K. Pandey, M. J. Reeves, C. J. Rodriguez, W. Rosamond, P. D. Sorlie, J. Stein, A. Towfighi, T. N. Turan, S. S. Virani, D. Woo, R. W. Yeh, M. B. Turner, A. H. A. S. Committee, S. S. Subcommittee, Heart Disease and Stroke Statistics-2016 Update: A Report From the American Heart Association. Circulation 133, e38–60 (2016).

2. W. H. Organization, Global status report on noncommunicable diseases 2014. World Health, 1–51 (2014).

3. A. M. Gotto, J. E. Moon, Pharmacotherapies for lipid modification: beyond the statins. Nature reviews. Cardiology 10, 560–570 (2013).

4. N. J. Stone, J. G. Robinson, A. H. Lichtenstein, C. N. Bairey Merz, C. B. Blum, R. H. Eckel, A. C. Goldberg, D. Gordon, D. Levy, D. M. Lloyd-Jones, P. McBride, J. S. Schwartz, S. T. Shero, S. C. Smith, K. Watson, P. W. F. Wilson, K. M. Eddleman, N. M. Jarrett, K. LaBresh, L. Nevo, J. Wnek, J. L. Anderson, J. L. Halperin, N. M. Albert, B. Bozkurt, R. G. Brindis, L. H. Curtis, D. DeMets, J. S. Hochman, R. J. Kovacs, E. M. Ohman, S. J. Pressler, F. W. Sellke, W.-K. Shen, S. C. Smith, G. F. Tomaselli, A. C. o. C. A. H. A. T. F. o. P. Guidelines, 2013 ACC/AHA guideline on the treatment of blood cholesterol to reduce atherosclerotic cardiovascular risk in adults: a report of the American College of Cardiology/American Heart Association Task Force on Practice Guidelines. Circulation 129, S 1–45 (2014).

5. G. K. Hansson, A.-K. L. Robertson, C. Söderberg-Nauclér, Inflammation and atherosclerosis. Annual review of pathology 1, 297–329 (2006).

6. P. Libby, Inflammation in atherosclerosis. Arteriosclerosis, thrombosis, and vascular biology 32, 2045–2051 (2012).

7. J. E. McLaren, D. R. Michael, T. G. Ashlin, D. P. Ramji, Cytokines, macrophage lipid metabolism and foam cells: implications for cardiovascular disease therapy. Progress in lipid research 50, 331–347 (2011).

8. K. J. Moore, I. Tabas, Macrophages in the pathogenesis of atherosclerosis. Cell 145, 341–355 (2011).

9. A. C. Calkin, P. Tontonoz, Transcriptional integration of metabolism by the nuclear sterol-activated receptors LXR and FXR. Nature reviews. Molecular cell biology 13, 213–224 (2012).

10. S.-S. Im, T. F. Osborne, Liver x receptors in atherosclerosis and inflammation. Circulation research 108, 996–1001 (2011).

11. N. Parikh, W. H. Frishman, Liver x receptors: a potential therapeutic target for modulating the atherosclerotic process. Cardiology in review 18, 269–274 (2010).

12. P. Costet, Y. Luo, N. Wang, A. R. Tall, Sterol-dependent transactivation of the ABC1 promoter by the liver X receptor/retinoid X receptor. The Journal of biological chemistry 275, 28240–28245 (2000).

13. A. Venkateswaran, B. A. Laffitte, S. B. Joseph, P. A. Mak, D. C. Wilpitz, P. A. Edwards, tP. Tontonoz, Control of cellular cholesterol efflux by the nuclear oxysterol receptor LXR alpha. Proceedings of the National Academy of Sciences of the United States of America 97, 12097–12102 (2000).

14. S. B. Joseph, A. Castrillo, B. a. Laffitte, D. J. Mangelsdorf, P. Tontonoz, Reciprocal regulation of inflammation and lipid metabolism by liver X receptors. Nature medicine 9, 213–219 (2003).

15. S. U. Naik, X. Wang, J. S. Da Silva, M. Jaye, C. H. Macphee, M. P. Reilly, J. T. Billheimer, G. H. Rothblat, D. J. Rader, Pharmacological activation of liver X receptors promotes reverse cholesterol transport in vivo. Circulation 113, 90–97 (2006).

16. J. J. Repa, S. D. Turley, J. A. Lobaccaro, J. Medina, L. Li, K. Lustig, B. Shan, R. A. Heyman, J. M. Dietschy, D. J. Mangelsdorf, Regulation of absorption and ABC1-mediated efflux of cholesterol by RXR heterodimers. Science (New York, N.Y.) 289, 1524–1529 (2000).

17. N. Levin, E. D. Bischoff, C. L. Daige, D. Thomas, C. T. Vu, R. A. Heyman, R. K. Tangirala, I. G. Schulman, Macrophage liver X receptor is required for antiatherogenic activity of LXR agonists. Arteriosclerosis, thrombosis, and vascular biology 25, 135–142 (2005).

18. R. K. Tangirala, E. D. Bischoff, S. B. Joseph, B. L. Wagner, R. Walczak, B. A. Laffitte, C. L. Daige, D. Thomas, R. A. Heyman, D. J. Mangelsdorf, X. Wang, A. J. Lusis, P. Tontonoz, I. G. Schulman, Identification of macrophage liver X receptors as inhibitors of atherosclerosis. Proceedings of the National Academy of Sciences of the United States of America 99, 11896–11901 (2002).

19. M. N. Bradley, C. Hong, M. Chen, S. B. Joseph, D. C. Wilpitz, X. Wang, A. J. Lusis, A. Collins, W. A. Hseuh, J. L. Collins, R. K. Tangirala, P. Tontonoz, Ligand activation of LXR beta reverses atherosclerosis and cellular cholesterol overload in mice lacking LXR alpha and apoE. The Journal of clinical investigation 117, 2337–2346 (2007).

20. S. B. Joseph, E. McKilligin, L. Pei, M. a. Watson, A. R. Collins, B. a. Laffitte, M. Chen, G. Noh, J. Goodman, G. N. Hagger, J. Tran, T. K. Tippin, X. Wang, A. J. Lusis, W. a. Hsueh, R. E. Law, J. L. Collins, T. M. Willson, P. Tontonoz, Synthetic LXR ligand inhibits the development of atherosclerosis in mice. Proceedings of the National Academy of Sciences of the United States of America 99, 7604–7609 (2002).

21. N. Terasaka, A. Hiroshima, T. Koieyama, N. Ubukata, Y. Morikawa, D. Nakai, T. Inaba, T-0901317, a synthetic liver X receptor ligand, inhibits development of atherosclerosis in LDL receptor-deficient mice. FEBS letters 536, 6–11 (2003).

22. A. Grefhorst, B. M. Elzinga, P. J. Voshol, T. Plösch, T. Kok, V. W. Bloks, F. H. van der Sluijs, L. M. Havekes, J. A. Romijn, H. J. Verkade, F. Kuipers, Stimulation of lipogenesis by pharmacological activation of the liver X receptor leads to production of large, triglyceride-rich very low density lipoprotein particles. The Journal of biological chemistry 277, 34182–34190 (2002).

23. J. R. Schultz, H. Tu, A. Luk, J. J. Repa, J. C. Medina, L. Li, S. Schwendner, S. Wang, M. Thoolen, D. J. Mangelsdorf, K. D. Lustig, B. Shan, Role of LXRs in control of lipogenesis. Genes & development 14, 2831–2838 (2000).

24. A. Kratzer, M. Buchebner, T. Pfeifer, T. M. Becker, G. Uray, M. Miyazaki, S. Miyazaki-Anzai, B. Ebner, P. G. Chandak, R. S. Kadam, E. Calayir, N. Rathke, H. Ahammer, B. Radovic, M. Trauner, G. Hoefler, U. B. Kompella, G. Fauler, M. Levi, S. Levak-Frank, G. M. Kostner, D. Kratky, Synthetic LXR agonist attenuates plaque formation in apoE-/-mice without inducing liver steatosis and hypertriglyceridemia. Journal of lipid research 50, 312–326 (2009).

25. D. Peng, R. Hiipakka, Q. Dai, J. Guo, Antiatherosclerotic effects of a novel synthetic tissue-selective steroidal liver X receptor agonist in low-density lipoprotein receptor-deficient mice. The Journal of Pharmacology and Experimental Therapeutics 22, 332–342 (2008).

26. T. Pfeifer, M. Buchebner, P. G. Chandak, J. Patankar, A. Kratzer, S. Obrowsky, G. N. Rechberger, R. S. Kadam, U. B. Kompella, G. M. Kostner, D. Kratky, S. Levak-Frank, Synthetic LXR agonist suppresses endogenous cholesterol biosynthesis and efficiently lowers plasma cholesterol. Current Pharmaceutical Biotechnology 12, 285–292 (2011).

27. E. M. Quinet, D. a. Savio, A. R. Halpern, L. Chen, C. P. Miller, P. Nambi, Gene-selective modulation by a synthetic oxysterol ligand of the liver X receptor. Journal of lipid research 45, 1929–1942 (2004).

28. B. A. Janowski, P. J. Willy, T. R. Devi, J. R. Falck, D. J. Mangelsdorf, An oxysterol signalling pathway mediated by the nuclear receptor LXR alpha. Nature 383, 728–731 (1996).

29. J. M. Lehmann, S. a. Kliewer, L. B. Moore, T. a. Smith-Oliver, B. B. Oliver, J. L. Su, S. S. Sundseth, D. a. Winegar, D. E. Blanchard, T. a. Spencer, T. M. Willson, Activation of the nuclear receptor LXR by oxysterols defines a new hormone response pathway. The Journal of biological chemistry 272, 3137–3140 (1997).

30. C. Yang, J. G. McDonald, A. Patel, Y. Zhang, M. Umetani, F. Xu, E. J. Westover, D. F. Covey, D. J. Mangelsdorf, J. C. Cohen, H. H. Hobbs, Sterol intermediates from cholesterol biosynthetic pathway as liver X receptor ligands. The Journal of biological chemistry 281, 27816–27826 (2006).

31. A. Radhakrishnan, L.-P. Sun, H. J. Kwon, M. S. Brown, J. L. Goldstein, Direct binding of cholesterol to the purified membrane region of SCAP: mechanism for a sterol-sensing domain. Molecular cell 15, 259–268 (2004).

32. M. S. Brown, J. L. Goldstein, Cholesterol feedback: from Schoenheimer′s bottle to Scap′s MELADL. Journal of lipid research 50 Suppl, S15–S27 (2009).

33. N. J. Spann, L. X. Garmire, J. G. McDonald, D. S. Myers, S. B. Milne, N. Shibata, D. Reichart, J. N. Fox, I. Shaked, D. Heudobler, C. R. H. Raetz, E. W. Wang, S. L. Kelly, M. C. Sullards, R. C. Murphy, A. H. Merrill, H. A. Brown, E. a. Dennis, A. C. Li, K. Ley, S. Tsimikas, E. Fahy, S. Subramaniam, O. Quehenberger, D. W. Russell, C. K. Glass, Regulated accumulation of desmosterol integrates macrophage lipid metabolism and inflammatory responses. Cell 151, 138–152 (2012).

34. J. Avigan, D. Steinberg, M. J. Thompson, E. Mosettig, Mechanism of Action of MER-29, An inhibitor of cholesterol biosynthesis. Biochemical and biophysical research communications 2, 63–65 (1960).

35. D. Steinberg, J. Avigan, E. B. Feigelson, Effects of Triparanol (Mer-29) on Cholesterol Biosynthesis and on Blood Sterol Levels in Man. The Journal of clinical investigation 40, 884–893 (1961).

36. T. J. Kirby, Cataracts produced by triparanol. (MER-29). Transactions of the American Ophthalmological Society 65, 494–543 (1967).

37. R. C. Laughlin, T. F. Carey, Cataracts in patients treated with triparanol. JAMA: the journal of the American Medical Association 181, 339–340 (1962).

38. R. K. Winkelmann, H. O. Perry, R. W. Achor, T. J. Kirby, Cutaneous syndromes produced as side effects of triparanol therapy. Archives of dermatology 87, 372–377 (1963).

39. C. P. Schaaf, J. Koster, P. Katsonis, L. Kratz, O. a. Shchelochkov, F. Scaglia, R. I. Kelley, O. Lichtarge, H. R. Waterham, M. Shinawi, Desmosterolosis-phenotypic and molecular characterization of a third case and review of the literature. American journal of medical genetics. Part A 155A, 1597–1604 (2011).

40. S. Yu, S. Li, A. Henke, E. D. Muse, B. Cheng, G. Welzel, A. K. Chatterjee, D. Wang, J. Roland, C. K. Glass, M. Tremblay, Dissociated sterol-based liver X receptor agonists as therapeutics for chronic inflammatory diseases. FASEB journal: official publication of the Federation of American Societies for Experimental Biology 30, 2570–2579 (2016).

41. E. G. Lund, L. B. Peterson, A. D. Adams, M.-H. N. Lam, C. A. Burton, J. Chin, Q. Guo, S. Huang, M. Latham, J. C. Lopez, J. G. Menke, D. P. Milot, L. J. Mitnaul, S. E. Rex-Rabe, R. L. Rosa, J. Y. Tian, S. D. Wright, C. P. Sparrow, Different roles of liver X receptor alpha and beta in lipid metabolism: effects of an alpha-selective and a dual agonist in mice deficient in each subtype. Biochemical pharmacology 71, 453–463 (2006).

42. E. M. Quinet, D. A. Savio, A. R. Halpern, L. Chen, G. U. Schuster, J.-A. Gustafsson, M. D. Basso, P. Nambi, Liver X receptor (LXR)-beta regulation in LXRalpha-deficient mice: implications for therapeutic targeting. Molecular pharmacology 70, 1340–1349 (2006).

43. T. G. Kirchgessner, P. Sleph, J. Ostrowski, J. Lupisella, C. S. Ryan, X. Liu, G. Fernando, D. Grimm, P. Shipkova, R. Zhang, R. Garcia, J. Zhu, A. He, H. Malone, R. Martin, K. Behnia, Z. Wang, Y. C. Barrett, R. J. Garmise, L. Yuan, J. Zhang, M. D. Gandhi, P. Wastall, T. Li, S. Du, L. Salvador, R. Mohan, G. H. Cantor, E. Kick, J. Lee, R. J. A. Frost, Beneficial and Adverse Effects of an LXR Agonist on Human Lipid and Lipoprotein Metabolism and Circulating Neutrophils. Cell metabolism 24, 223–233 (2016).

44. H. C. Andersson, L. Kratz, R. Kelley, Desmosterolosis presenting with multiple congenital anomalies and profound developmental delay. American Journal of Medical Genetics 113, 315–319 (2002).

45. D. R. FitzPatrick, J. W. Keeling, M. J. Evans, a. E. Kan, J. E. Bell, M. E. Porteous, K. Mills, R. M. Winter, P. T. Clayton, Clinical phenotype of desmosterolosis. American journal of medical genetics 75, 145–152 (1998).

46. W. Chen, G. Chen, D. L. Head, D. J. Mangelsdorf, D. W. Russell, Enzymatic reduction of oxysterols impairs LXR signaling in cultured cells and the livers of mice. Cell metabolism 5, 73–79 (2007).

47. E. Seki, S. De Minicis, C. H. Osterreicher, J. Kluwe, Y. Osawa, D. A. Brenner, R. F. Schwabe, TLR4 enhances TGF-beta signaling and hepatic fibrosis. Nat Med 13, 1324–1332 (2007).

48. I. Mederacke, D. H. Dapito, S. Affo, H. Uchinami, R. F. Schwabe, High-yield and high-purity isolation of hepatic stellate cells from normal and fibrotic mouse livers. Nat Protoc 10, 305–315 (2015).

